# Visual quality control with *CytoMDS*, a Bioconductor package for low dimensional representation of cytometry sample distances

**DOI:** 10.1101/2024.07.01.601465

**Authors:** Philippe Hauchamps, Simon Delandre, Stéphane T. Temmerman, Dan Lin, Laurent Gatto

**Author notes:** Computational Biology and Bioinformatics, de Duve Institute, UCLouvain, Av. Hippocrate 75, 1200 Brussels, Belgium. **Data availability:** Raw cytometry data files are available either on Zenodo (DOI:10.5281/zenodo.10572228) or FlowRepository (ID: FR-FCM-ZYL8). The *ImmunoSenescence Human PBMC* dataset is available upon request. **Code availability:** All code needed to reproduce the results presented in the current article is available on the following GitHub repository: github.com/UCLouvain-CBIO/2024-CytoMDS-code. **Funding:** This work was funded by GlaxoSmithKline Biologicals S.A., under a cooperative research and development agreement between GlaxoSmithKline Biologicals S.A. and de Duve Institute (UCLouvain). **Competing interests:** S.D., S.T., and D.L. are employees of the GSK group of companies, and report ownership of GSK shares. S.T. is listed as inventor on patents owned by the GSK group of companies. P.H. is a student at the de Duve Institute (UCLouvain) and participates in a post graduate studentship program at GSK.

## Abstract

Quality Control (QC) of samples is an essential preliminary step in cytometry data analysis. Notably, identification of potential batch effects and outlying samples is paramount, to avoid mistaking these effects for true biological signal in downstream analyses. However, this task can prove to be delicate and tedious, especially for datasets with dozens of samples.

Here, we present *CytoMDS*, a Bioconductor package implementing a dedicated method for low dimensional representation of cytometry samples composed of marker expressions for up to millions of single cells. This method allows a global representation of all samples of a study, with one single point per sample, in such a way that projected distances can be visually interpreted. *CytoMDS* uses *Earth Mover’s Distance* for assessing dissimilarities between multi-dimensional distributions of marker expression, and *Multi Dimensional Scaling* for low dimensional projection of distances. Some additional visualization tools, both for projection quality diagnosis and for user interpretation of the projection coordinates, are also provided in the package.

We demonstrate the strengths and advantages of *CytoMDS* for QC of cytometry data on three real biological datasets, revealing the presence of low quality samples, batch effects and biological signal between sample groups.

## Introduction

Recent technology advances in flow and mass cytometry allow the acquisition of high dimensional sample data, with up to respectively 40 and 50 parameters measured simultaneously for millions of cells (Kare et al., 2023, Spitzer and Nolan, 2016). For such high dimensional data samples, extracting relevant biological information becomes challenging (Newell and Cheng, 2016). Moreover, in large clinical studies, possibly with some longitudinal and multicentric components, this difficult task is itself multiplied by the high number of samples (Hartmann et al., 2019, Schmit, Klomp, and Khan, 2021). In the last decade, this led to the development of (semi) automated data analysis pipelines (Saeys, Van Gassen, and Lambrecht, 2016), to complement the traditional manual gating approaches (Staats et al., 2019). Among the different steps involved in (automated) data analysis pipelines, pre-processing and Quality Control (QC) of data samples is paramount, as it guarantees that the downstream analyses are not compromised by confounding noise. For cytometry data, pre-processing and QC involve different treatments and checks (Liechti et al., 2021): compensation, stability of signal in time, removal of undesirable events like debris, doublets or dead cells, batch effect detection and removal, etc.

Whatever the level of automation of these treatments, some manual checks of the cleaned samples, possibly at different stages of the pre-processing, are still necessary. This is typically done by human operators using interactive visual software, like e.g. FlowJo or Cytobank - among the available commercial platforms, or free open source solutions (Hammill, 2021, Hauchamps et al., 2024). Nevertheless, these tools are mostly designed to check samples one by one, or possibly to visually compare two or a handful of samples at once. However, two categories of QC, namely outlying sample identification (Chen et al., 2020) and batch effect detection (Goh, Wang, and Wong, 2017), would benefit from a global representation of the dataset. Well-known classical data exploration techniques like *Principal Component Analysis (PCA)* (Pearson, 1901) have the potential to provide such a global view of the data, and are routinely used in other omics context like e.g. bulk transcriptomics (Chen et al., 2020) and bulk proteomics (Goh and Wong, 2017) data analyses. In bulk experiments, each sample results in a single vector of measurements because all its cells are merged. All samples can be combined into a single matrix which corresponds to the use of a standard PCA, or a similar algorithm. In single cell experiments, there is a measurement vector for each cell and each sample results in a measurement matrix. Applying these bulk techniques to cytometry data presents a major challenge in summarising each sample by a vector for comparing samples (not their cells) with each other.

To address this challenge, we developed *CytoMDS* (Hauchamps and Gatto, 2024), an R package for low dimensional projection of cytometry samples, which allows a global representation of all samples of a study, with one single point per sample, in such a way that projected distances can be visually interpreted. In this article, we present the *CytoMDS* method, and demonstrate its use on three real cytometry datasets, illustrating the detection of various quality issues. Finally we compare our approach with similar existing methods for global problem detection in cytometry samples, and discuss its advantages and potential limitations. *CytoMDS* is available in Bioconductor (Huber et al., 2015), as of version 3.19.

## Materials and Methods

### Proposed method

Figure 1 sketches an overview of the *CytoMDS* workflow. As input, the method requires a set of minimally pre-processed - i.e. scale transformed and, if applicable, compensated or unmixed - sample files in *flow cytometry standard (fcs)* format (Figure 1, [A]). The *CytoMDS* method itself (Figure 1, [B]) can be decomposed into two main steps:

**Figure 1.**
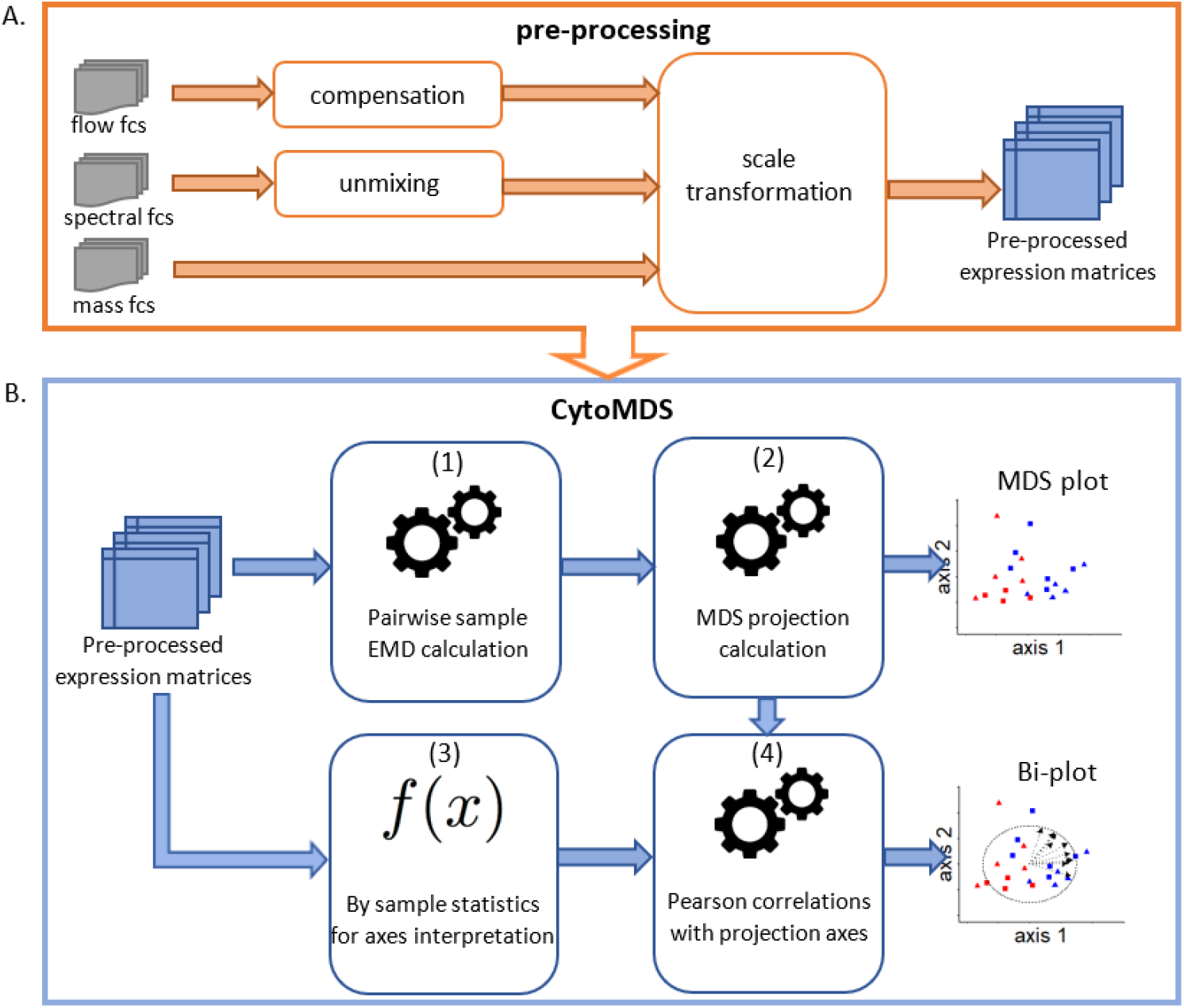
Overview of the *CytoMDS* workflow. Pre-processing of the raw *fcs* files [A] should be done according to the workflow in the orange box, which includes scale transformation, and might include compensation or unmixing depending on the type of data. The *CytoMDS* method itself [B] takes as input one pre-processed expression matrix per sample. First, *Earth Mover’s Distances (EMD)* are calculated for each pair of samples (1). Second, the resulting distance matrix is projected using *Multi Dimensional Scaling (MDS)* (2), producing a *MDS* plot. Optionally, in order to interpret *MDS* coordinates and plot directions, *CytoMDS* can compute a set of chosen sample statistics (3). *CytoMDS* then calculates Pearson correlations of these statistics with the *MDS* coordinates (4), producing one or several bi-plot(s).

- computing the distances between each sample pair (1);
- projecting the resulting pairwise distance matrix onto a low dimensional space (2) of dimension *p, p* being typically 2, 3 or a small number sufficient to reach a good projection quality (see below).

As a result, each sample is a single point in this *p*-dimensional space, and all samples can be represented in plots combining those *p* dimensions. Optionally, *bi-plots* can be performed by overlaying on these plots the correlations (4) of chosen sample statistics (3) with the projection coordinates, in order to interpret the directions within the plots.

#### Distance calculation

In order to characterize the dissimilarity between samples, the *Earth Mover’s Distance (EMD)*, a.k.a. the *Wasserstein distance of order one*, is chosen as our approach. Considering two samples *a* and *b*, which are represented by their respective *M*-dimensional distribution of expression intensities for the *M* markers, the *EMD* can be seen as the minimal cost (or effort) needed for transporting the distribution mass of one sample to the other, where the cost of transporting one unit of mass from location *x* to location *y* is given by the Euclidean distance between *x* and *y* (Rubner, Tomasi, and Guibas, 2000).

The *EMD* has been shown to be suitable for quantifying dissimilarities between cytometry samples, and to provide better insight than other types of metrics for this purpose (Orlova et al., 2016). It has been used in different applications of cytometry data analysis, especially in the evaluation of batch correction methods (Van Gassen et al., 2020, Pedersen et al., 2022) or for obtaining a sample representation in relation with different phenotypes (Yi and Stanley, 2022).

Here, for computational efficiency given the potentially high number of markers, we approximate the multi-dimensional *EMD*, by the sum of the *M* 1D *EMD* for each marker *m*: 1,… *M*. Assuming that each unidimensional marker expression distribution has been discretized in 1D histograms consisting of a number of bins with specific location and weight, the 1D *EMD* is then formally defined as follows:

- Let *a*_*i*_ and 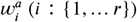 be respectively the location and weight assigned to bin *i* of the 1D histogram of marker *m* in sample *a*.
- Let *b*_*j*_ and 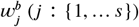 be respectively the location and weight assigned to bin *j* of the 1D histogram of marker *m* in sample *b*.

The empirical distribution functions *F*(*t*) and *G*(*t*) of the marker *m* intensity, respectively for sample *a* and *b*, are therefore: 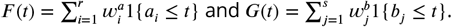

The 1D *EMD* between marker *m* intensities of samples *a* and *b* is then:

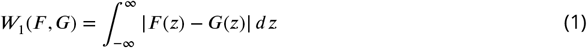

To compute the *EMD* for each marker (Equation 1), we take advantage of the R package *transport* (Schuhmacher et al., 2024). This computation is then repeated for each pair of samples, leading to a symmetric pairwise distance matrix having *N* rows and *N* columns - with *N* the number of samples. Its diagonal is filled with zeros.

#### Projection algorithm

The second step consists of projecting the distance matrix of dimensions *NxN* onto a low dimensional embedding with *p* dimensions. As our goal is to visualize sample distances to identify outliers or batch effects, we chose a method from the *Multi Dimensional Scaling (MDS)* family, since they are good at preserving distances and the global data structure (Lee and Verleysen, 2007), in contrast to popular local neighbourhood preserving methods such as *t-SNE* (van der Maaten and Hinton, 2008) and *UMAP* (McInnes, Healy, and Melville, 2018).

Furthermore, because the *EMD* is a metric but is not euclidean, we need to resort, for the projection, to *non classical metric MDS*. It is a non linear method minimizing a specific criterion called *stress* (Lee and Verleysen, 2007) that represent a quadratic error of projection and is defined as follows:

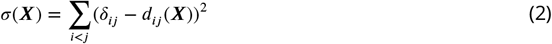

where ***X*** is the low-dimensional embedding, *δ*_*ij*_ is the original *EMD* between sample *i* and sample *j*, and *d*_*ij*_ is the euclidean distance between the corresponding projections in ***X***.

In order to minimize the stress criterion (Equation 2), *CytoMDS* uses the *SMACOF* (Scaling by MAjorizing a COmplicated Function) algorithm, implemented in the *smacof* R package (de Leeuw and Mair, 2009). Finally, the projection axes of the *p*-dimensional embedding are linearly transformed using *PCA*. This last step results in new projection coordinates that are ordered in decreasing percentages of variance, as in a classical *PCA* visualization. However, here these percentages are with respect to the total variance associated with the embedding, which differs from the total variability of the data.

#### Quality of projection

*CytoMDS* provides different diagnostic tools to assess the quality of the low dimensional projection. The first one is a visualization plot, called *Shepard’s diagram*, which allows to assess how close the projected euclidean distances are to their *EMD*, for each sample pair. Examples of such diagrams can be found in Figure S1 and Figure S3. The second diagnostic tool is a global indicator of the projection quality, the *pseudoR*^2^, which assesses to which extend the pairwise sample *EMD* can be explained by the low dimensional euclidean distances on the projection. It is defined as follows:

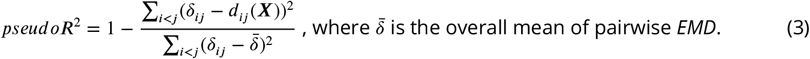

This *pseudoR*^2^ plays a central role in *CytoMDS* as, by default, *p*, the number of dimensions of the low dimensional embedding, is tuned automatically to reach a specific *pseudoR*^2^ target for the projection, 0.95 by default. Finally, additional indicators from the *SMACOF* algorithm can also be recovered, such as the minimised value of the stress criterion (Equation 2) and the contribution of each sample to the stress, i.e. the *stress per point* (de Leeuw and Mair, 2009).

#### Coordinates and directions interpretation using bi-plots

*Bi-plots* are plots that are overlaid to the *MDS* projection, allowing to interpret directions of interest by relating them to well-chosen sample characteristics (Borg, Groenen, and Mair, 2018). Creating a bi-plot involves choosing *external scales*, i.e. sample specific characteristics that can be fitted to the *MDS* space. Possible choices are summary statistics of marker intensities, e.g. univariate mean, standard deviation, median or other quantiles, etc., or any other sample representative characteristics (multivariate statistics, number of events in sample, etc.). *CytoMDS* calculates the Pearson correlations of the *external scales* with respect to the visualized *MDS* coordinates, and overlays arrows on the plot, one for each *external scale*, having coordinates equal to their respective correlations. A unit 1 correlation circle is also plotted as landmark for visual convenience (e.g. in Figure 2, [C] and [D]).

**Figure 2.**
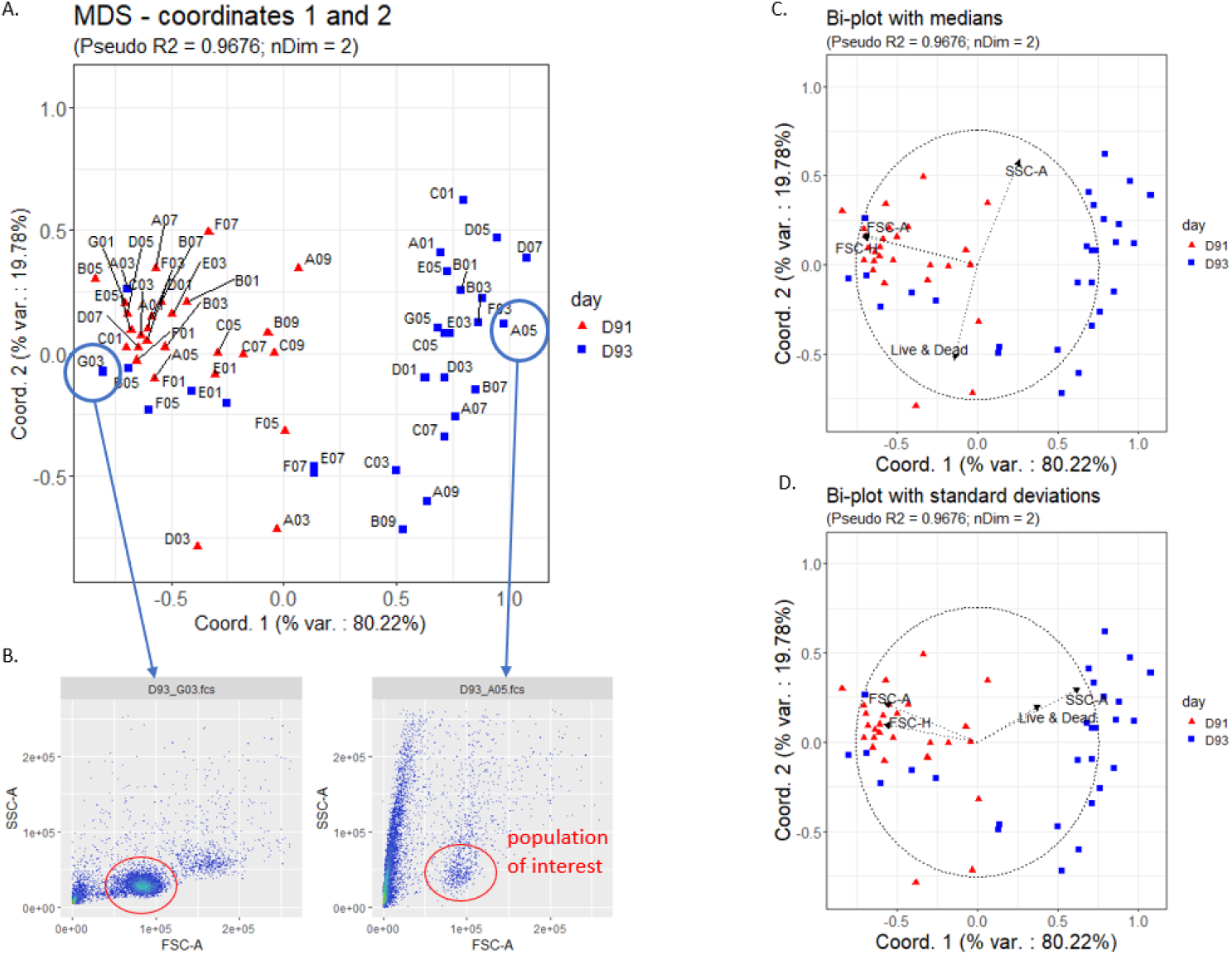
The low dimensional projection of the *HBV Chronic Mouse* dataset [A] reveals a clear split between two sample groups, not fully attributable to a batch effect linked to day of acquisition - since a minority of day D93 samples belong to the left sample group. Bi-plots overlaying respectively channels medians [C], and standard deviations [D] show that the x axis, which is the direction discriminating the two sample groups, is strongly negatively correlated both with *FSC* channel median and *FSC* standard deviation, and also positively correlated with *SSC* standard deviation. A comparison of the *FSC* vs. *SSC* 2D representation of selected samples having opposite x coordinates [B] shows that the right sample group contains low quality samples, with very few events belonging to the cell population of interest.

### Example datasets

In order to demonstrate *CytoMDS* in subsequent sections, we make use of three datasets.

#### HBV Chronic Mouse dataset

The *HBV Chronic Mouse* dataset is a flow cytometry dataset, collected in a preclinical study aimed at assessing the effect of different therapeutic vaccine regimes on the immune response of Hepatitis B Virus transduced mice. This dataset has been described in Hauchamps et al., 2024. In short, it contains 55 flow cytometry mouse liver lymphocyte samples, analyzed with a 12-channel flow cytometry panel. The 55 mice were split in 5 experimental groups corresponding to different vaccine regimens, and the flow cytometry data were acquired in two flow cytometry batches performed on two different days (D91 and D93). The *HBV Chronic Mouse* dataset is available on Zenodo (DOI:10.5281/zenodo.8425840).

#### ImmunoSenescence Human PBMC dataset

The *ImmunoSenescence Human PBMC* dataset is also a flow cytometry dataset. This dataset has been generated in the context of the development of a 26 markers panel that aimed at phenotyping all the main immune cell populations present in PBMC (Peripheral Blood Mononuclear Cell) samples. This readout development was part of a study that aimed at assessing the effect of age on the human immune system. The dataset contains 20 *fcs* files, corresponding to the flow cytometry data acquisition of 15 PBMC samples. These were collected and cryopreserved from healthy individuals, split into two age groups - younger and older adults - and were part of two different GSK biobanks. There is therefore a potential confounding effect between the biological signal of interest - linked to ageing - and the two different sample origins. The flow cytometry data acquisition itself was performed in two different acquisition rounds (a *former* and a *later* data acquisition), and 5 of the biological samples were acquired in both rounds (see Table S1). Sampling and research were approved by the Belgian Ethics committee and written informed consent was obtained from all participants prior to sample collection, in accordance with the Helsinki Declaration.

#### Krieg_Anti_PD1 dataset

The *Krieg_Anti_PD1* dataset is a mass cytometry dataset (Krieg et al., 2018), consisting of 20 baseline samples (prior to treatment) of peripheral blood from melanoma skin cancer patients subsequently treated with anti-PD-1 immunotherapy. The samples were split across 2 conditions (non-responders and responders) and in 2 acquisition batches. The goal of the study was to identify biomarkers of responsiveness to immunotherapy at baseline. This dataset is available in FlowRepository (ID: FR-FCM-ZYL8) and is directly accessible in R from the Bioconductor data package *HDCytoData* (Weber and Soneson, 2019).

## Results

### Identifying low quality samples

In this section, we demonstrate the use of *CytoMDS* on the *HBV Chronic Mouse* dataset, with QC as a specific goal. The raw data were pre-processed by applying compensation and scale transformation using respectively bi-exponential transformations (Parks, Roederer, and Moore, 2006) for fluorescent channels, and linear transformations for *FSC* and *SSC* channels. Coefficients of the linear transformations were calibrated as to align 5 and 95 quantiles of *FSC/SSC* channels to those of a reference fluorescent channel. *CytoMDS* was used to compute *EMD* between each pair of samples. Here, since the focus is on tracking sample quality issues, we limited ourselves to four specific channels as input of the *EMD* calculation, i.e. *FSC-A, FSC-H, SSC-A* and the *Live/Dead* channels, which are the channels used for pre-gating (den Braanker, Bongenaar, and Lubberts, 2021). Finally the low dimensional projection was computed for the 55 samples of the dataset.

The *Shepard’s diagram*, in Figure S1, illustrates the projection quality. It shows that most pairwise *EMD* are close to the 45 degree identity line. Also, the *pseudo R*^2^ is 0.9679, hence above the 0.95 threshold (chosen by default), with only two projected dimensions.

The projection, in Figure 2 [A], highlights two major clusters of samples. On the positive side of the x coordinate, we can find most of the samples acquired on day D93 (blue squares). The other cluster of samples, on the negative side of the x coordinate, includes mostly the samples from day D91 (red triangles), but also a minority of day D93 samples. Therefore, these clusters cannot be attributed solely to a batch effect of acquisition time point. Two bi-plots using medians and standard deviations (Figure 2, respectively [C] and [D]) show that the x coordinate is strongly correlated, in a negative manner, with both the *FSC-A* channel median and *FSC-A* standard deviation, and positively correlated with *SSC-A* standard deviation. *FSC-A* and *SSC-A* are therefore taken as plot axes in Figure 2 [B], showing 2D plots for two representative samples, selected for their extreme opposite position on the first axis of the projection. Comparing these two plots reveals that sample *D93_A05* is a low quality sample where the cell population of interest - liver lymphocytes - is extremely small, and where most events seem to correspond to debris or dead/dying cells, whereas sample *D93_G03* is a good quality sample. Actually, as shown in Figure S2, the same pattern can be seen in all samples belonging to the low quality cluster.

As a conclusion, the *CytoMDS* projection of the *HVB Chronic Mouse* dataset, complemented by biplots, has allowed to identify a sample cluster corresponding to low quality samples, in terms of relative percentage of events belonging to the cell population of interest.

### Highlighting batch effects

In this section, we apply *CytoMDS* on the *ImmunoSenescence Human PBMC* dataset. As in the previous section, we pre-processed the data by applying compensation - based on a manually adjusted compensation matrix - and scale transformed the channel intensities - using bi-exponential transformations or linear transformations as in previous section. The pairwise sample *EMD* were calculated using the *FSC-A, FSC-H, SSC-A* and *Live/Dead* channels. In contrast to the *HBV Chronic Mouse* dataset, *CytoMDS* required 3 dimensions to reach the *pseudo R*^2^ quality threshold of 0.95. The *Shepard’s diagram* is shown on Figure S3.

Figure 3 shows the low dimensional projection according to 2 different combinations of *MDS* axes. Figure 3 [A] shows the projection on axes 1 and 2, which reveals a clear separation between the samples of the former (triangles), and later (squares) data acquisitions. This is an obvious batch effect. Using bi-plots to interpret further the direction of separation highlights that this batch effect is almost exclusively explained by a difference of location and scale of the *Live/Dead* channel (Figure S4). This effect is most probably caused by technical differences between the two data acquisitions for this particular channel. These could be, for example, due to different cytometer set-ups, or different staining efficiencies for the different batches.

**Figure 3.**
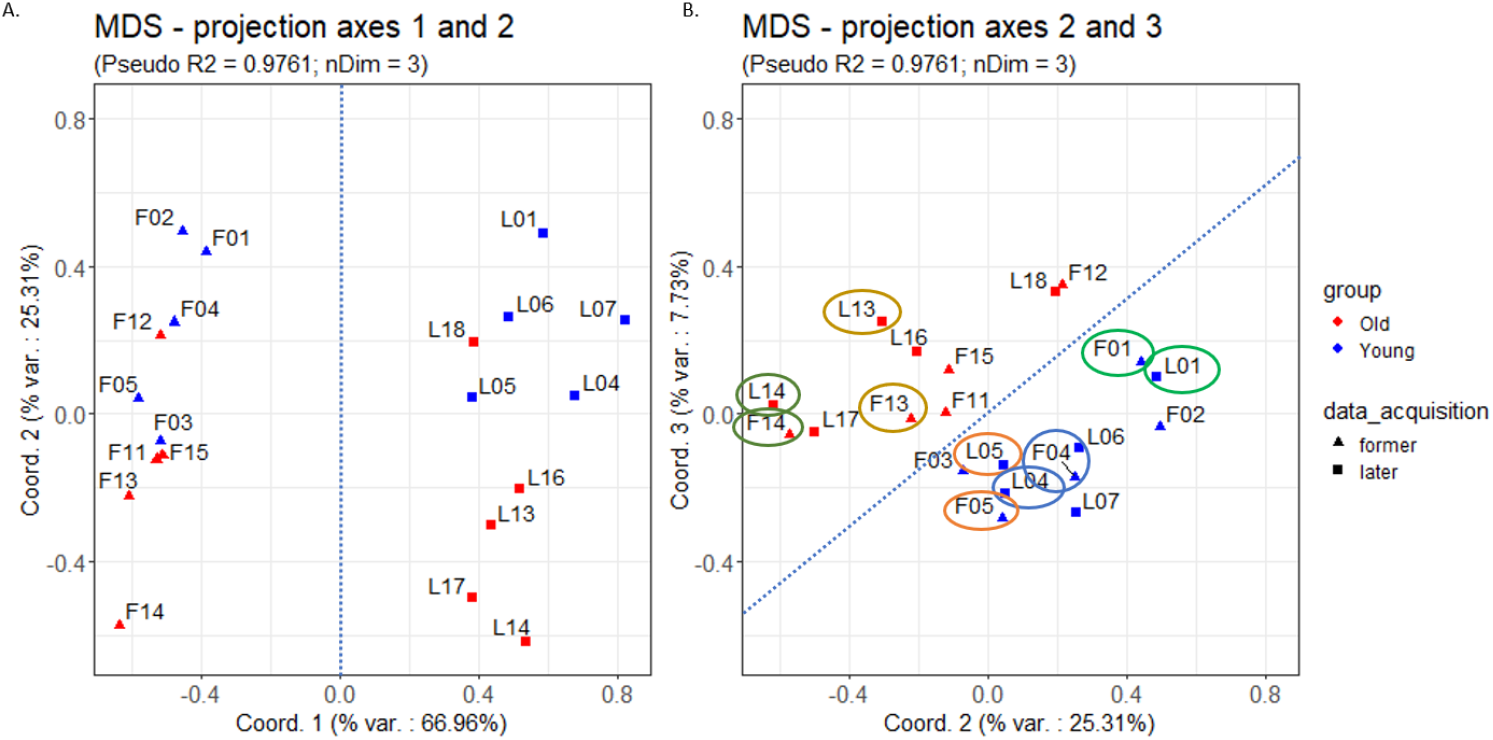
Low dimensional projection of the *ImmunoSenescence Human PBMC* dataset. (A) The projection on axes 1 and 2 reveals a clear batch effect between data points belonging to the former (triangles) and the later data acquisitions (squares). (B) The two groups (‘old’ in red vs. ‘young’ in blue), can be separated along a direction that is a linear combination of axes 2 and 3. On the same plot, one can also notice the proximity of the data points corresponding to the biological samples from the same donor (ellipses of different colours: F01 vs. L01, F04 vs. L04, F05 vs. L05, F13 vs. L13, F14 vs. L14).

On Figure 3 [B], samples of the 2 experimental groups (old vs. young donors) can be separated along a direction that is a linear combination of axes 2 and 3. Since we have only included pre-processing channels in the distance calculation, this separation is unlikely to reflect the biological signal of interest. In fact, it probably highlights another batch effect, which is confounded with experimental groups. We show this by interpretation of the direction of separation, which is here less obvious. Bi-plots using medians (Figure S5 [C]), or using standard deviations (not shown), are not able to identify any strong correlation with the direction of separation. However, Figure S5 [D] shows that the latter is strongly correlated with 10th quantiles of *FSC* and *SSC* channels, highlighting the tendency for young adult samples to contain more debris than old adult samples (Figure S5 [B]). This is most probably due to differences in sample handling between these two groups of samples, which were taken from different biobanks (see Methods). Finally, one can also notice that, apart from the first coordinate, driven by *Live/Dead* channel intensities, sample data points with the same label number tend to colocate (ellipses on Figure 3 [B]). This reflects the fact that these pairs correspond to two acquisitions of the same original biological sample.

To summarize this section, the *CytoMDS* projection of the *ImmunoSenescence Human PBMC* dataset, according to two different combinations of axes, allowed to highlight separations of sample groups that could be attributed to two different batch effects, respectively linked to data acquisition time point and sample origin. Besides, proximity of the samples corresponding to the same biological donor could be shown by the projection.

### Highlighting biological signal between sample groups

*CytoMDS* projections can also reveal biological signal between different groups of samples, provided that this signal is sufficiently strong. In this section, we apply *CytoMDS* on the *Krieg_Anti_PD1* dataset. As this dataset is a mass cytometry dataset, pre-processing involves applying an *arcsinh()* scale transformation, but no channel compensation. Here, we used all 25 channels as input for the *EMD* computation, and projected the pairwise distances between the 20 samples using *CytoMDS* with 2 dimensions (Figure 4, [A]).

**Figure 4.**
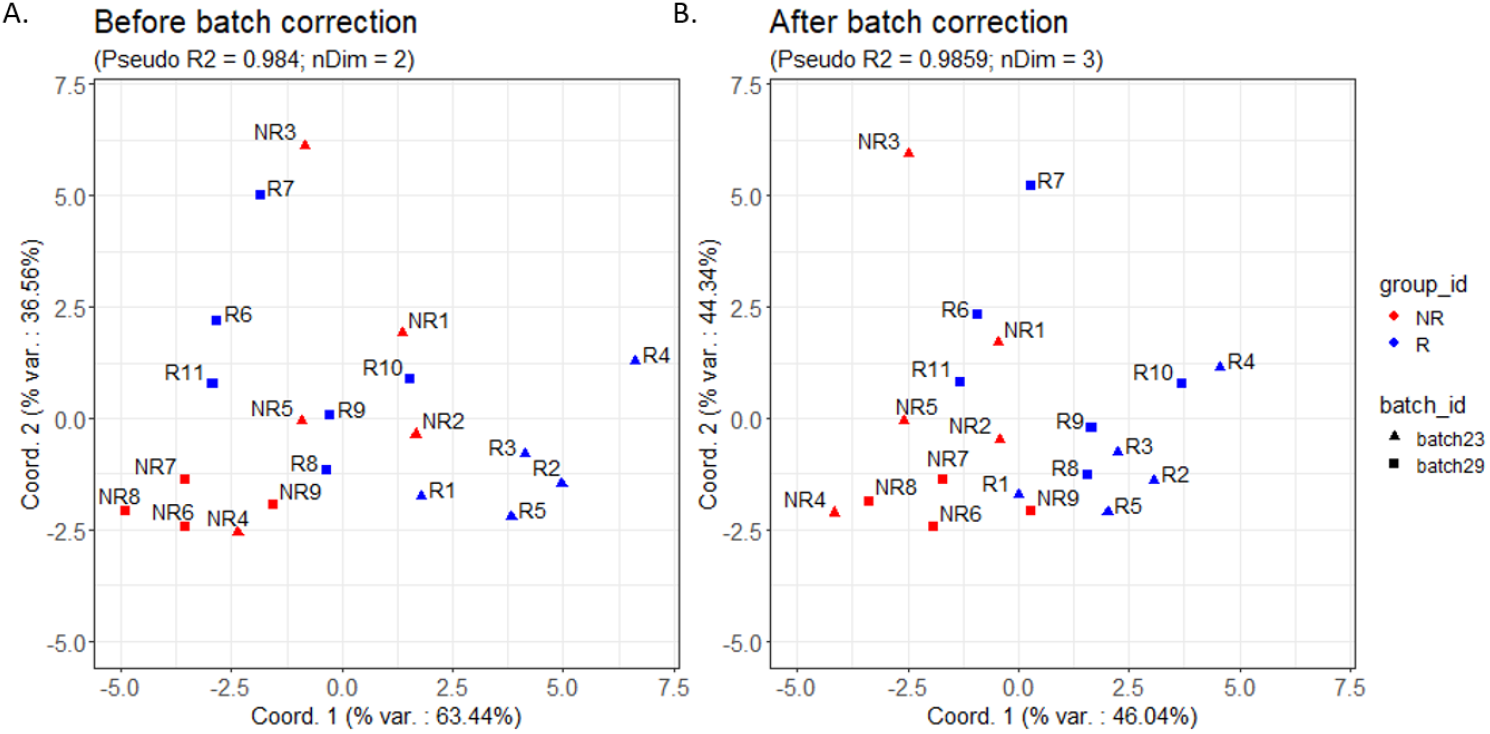
Low dimensional projection of the *Krieg_Anti_PD1* dataset, before batch correction [A], and after batch correction [B], on the first two axes. [A] The projection reveals a clear batch effect between *batch29* samples (squares), which tend to be located on the left side of the plot, and the *batch23* samples (triangles), which tend to be located on the right side. [B] The batch correction results in a better mix between samples of different batches. As a consequence, the separation between the two condition groups - responders (blue) vs. non responders (red) - is more sensible.

The *Krieg_Anti_PD1* dataset contains a strong batch effect, due to sample acquisition on two different days (Krieg et al., 2018). This batch effect is clearly visible on Figure 4, [A]: *batch29* samples (squares) tend to be located on the left of the plot, while the *batch23* samples (triangles) tend to be located on the right. Therefore, we performed a batch correction to this dataset, by applying the *ComBat* method (Johnson, Li, and Rabinovic, 2007), implemented in the *cyCombine* package (Pedersen et al., 2022), and using the responder status - on anti-PD-1 immunotherapy - as covariate. Figure 4 [B] displays the projection after batch correction, showing a more sensible separation of the two biological condition groups - responders (blue) vs. non responders (red) - and a better mixing of samples from the two batches.

Figure S6 shows bi-plots with medians, for the projection on axes 1 and 2, after batch correction. In the original study (Krieg et al., 2018), it was highlighted that the frequency of a small subpopulation of monocytes in baseline samples was positively associated with responder status following immunotherapy treatment. The left bi-plot of Figure S6 shows that the 11 markers involved in the definition of this rare population tend to be positively correlated with the direction of separation between responders vs non responders, while this is not the case on the right bi-plot, which shows the other 14 markers.

As a conclusion of this section, we showed that *CytoMDS* can be used, as an exploratory data visualization tool, to highlight biological signal between groups of samples, provided this signal is sufficiently strong.

## Discussion

### *CytoMDS* enables fast and global detection of data quality issues

In the Results section, we have provided three examples of using *CytoMDS* to detect sample quality issues. For the *HBV Chronic Mouse* dataset, the issue was specific to a group of samples for which an improper sample handling problem had shrunk the population of interest to a small percentage of the acquired events. For the *ImmunoSenescence Human PBMC* dataset and the *Krieg_Anti_PD1* dataset, the issues were the presence of one or several batch effects, sometimes confounded with the biological condition of interest. Beyond the identification of these quality issues, the ability to link these to specific sample characteristics, hence to potential root causes, is essential to the users, who will more easily make educated decisions about the encountered problems. For this purpose, the *CytoMDS* method has several strengths:

- It generates a global view of all samples, where the distances between points can be quantitatively interpreted.
- It provides bi-plots, a powerful interpretation tool which enables to correlate plot coordinates to sample characteristics. As a benefit, valuable insight on the potential cause(s) of the observed pattern(s) can be inferred.
- It allows unexpected batch effects to be revealed, in addition to those suspected. Indeed, it is not necessary to know in advance which samples belong to which batch. The *MDS* projection itself can be used to uncover patterns, which can in turn be dissected with the help of bi-plots, in order to highlight the specific sample characteristics that explain the observed groupings of samples. Similarly, this process can be used to detect outlier samples.

These strengths are not found in tSNE or UMAP, which are other popular techniques used to highlight batch effects, and which typically project the events of all samples of a dataset at once with low dimensions. An example of a 2 dimensional tSNE plot of the *HBV Chronic Mouse* dataset is provided in Figure S7. This plot suggests a batch effect driven by acquisition day, while we know from Figure 2 that this effect is driven by a quality issue observed for only a subset of the samples acquired on a specific day. However, as is shown in Figure S7, due to the high number of samples and high number of cells, there is no easy way to precisely identify the impacted samples, nor to link the quality issue to specific sample characteristics. In contrast, the *CytoMDS* projection (Figure 2) stays informative when the number of samples increases to tens or even a few hundreds.

Nowicka et al., 2017 proposed another *MDS* projection method for QC of cytometry data. This method, however, is not based on distributional distances between samples, but uses the medians of each channel to summarize the information contained in each sample. As a result, this method can lead to projections that are very similar to the *CytoMDS* ones, only when sample distances are driven mostly by the channel medians, but is insensitive to other differences between sample distributions. Figure S8 shows a comparative example, with low dimensional projections of the *ImmunoSenescence Human PBMC* dataset. *CytoMDS* allows to identify the distributional charcateristics that drive the dissimilarity between the samples at hand, which are not necessarily the channel medians.

### *CytoMDS* can be used at different stages of the data analysis pipeline

In Figure 1, we proposed a workflow where only compensation and scale transformation were performed prior to *CytoMDS* projection. However, as a data exploration tool, *CytoMDS* can be run at any stage of the data analysis pipeline. In the Results section, the *CytoMDS* projections were compared before and after batch correction. In general, the iterative use of *CytoMDS* can uncover quality issues at any stage of the data analysis pipeline.

### Computation time and scalability to larger datasets

In the Supplementary Results, we present a benchmarking of the *CytoMDS EMD* computation time. As is shown in Figures S9 and S10, the number of markers impacts the *EMD* computation time linearly, while sub-sampling allows to mitigate the effect of a large number of events per sample, with no material impact on precision. As a result, a single *EMD* calculation requires less than one second.

However, for datasets with hundreds of cytometry samples, computing the *EMD* between all sample pairs is a heavy computational task. A first pitfall might arise when loading the whole dataset in memory, which might not be possible due to its size. Another challenge is that calculating a matrix of pairwise distances has a computational complexity of *O*(*N*^2^), which can lead to long computation time for large datasets. The *CytoMDS* package provides two mechanisms to mitigate these issues. First, to handle datasets of greater size than the available computer RAM, *CytoMDS* provides a distance calculation method that dynamically loads data on the fly when calculating a pairwise sample distance, and releases the corresponding memory afterwards. Second, the user can instruct *CytoMDS* to parallelize the distance calculation process. In that case, the calculation engine takes advantage of the *BiocParallel* package (Morgan et al., 2023) to dispatch the calculation on several workers. The calculation engine then automatically creates balanced worker tasks, corresponding to blocks of the distance matrix, minimizing the overlap between data loaded in memory by different workers. These mechanisms have allowed to decrease the computation time of the full pairwise distance matrix of a dataset of 1400 mass cytometry samples with 43 parameters, from about 7 days to about 1.3 hours on a 14 cores user laptop (Granjeaud et al., 2024).

### Positioning, limitation and possible future extension

*CytoMDS* has been developed as an exploratory visualization method, aiming at facilitating the quality control process, for outliers and batch effects detection. As was shown in the *Krieg_Anti_PD1* example, this exploratory visualization can also highlight sample clusters that correspond to different conditions of interest, i.e. pointing to the existence of a biological signal. However, identification of such signal requires a dedicated differential expression/abundance analysis method (e.g. Weber et al., 2019, Seiler et al., 2021). In particular, a biological signal of interest consisting in the existence (or absence) of a rare immune cell population can translate into very subtle sample distribution differences involving conditional distributions of multiple combined marker intensities. These cannot be captured by *EMD* calculated as sum of marginal distributions of markers, as is done in *CytoMDS* (see Methods). A possible extension of this work could be to refine the calculation of pairwise sample *EMDs*, to incorporate the information on joint marker distributions. This would however involves a difficult computational challenge, especially in high dimensional cytometry datasets. Finally, the current implementation has already shown, throughout the different real dataset examples, its ability to highlight various data quality issues, and link them to potential root causes, which is the main goal of our method.

## Declarations

### Author contributions

P.H.: conceptualization (lead); methodology; software (lead); writing - original draft (lead). S.D.: methodology (supporting); writing - original draft (supporting). S.T.: supervision (supporting); writing - review & editing (supporting). D.L.: supervision; methodology (supporting); writing - review & editing. L.G.: supervision; conceptualization (supporting); methodology; software (supporting); writing - review & editing.

### Availability of data and materials

Raw cytometry data files are available either on Zenodo (DOI:10.5281/zenodo.10572228) or FlowRepository (ID: FR-FCM-ZYL8). The *ImmunoSenescence Human PBMC* dataset is available upon request.

### Code availability

*CytoMDS* package is available on Bioconductor: https://www.bioconductor.org/packages/release/bioc/html/CytoMDS.html

All code needed to reproduce the results presented in the current article is available on the following GitHub repository: https://github.com/UCLouvain-CBIO/2024-CytoMDS-code.

## Acknowledgments

The authors would like to thank Mehdi Hamrouni and Babak Bayat (GSK) for the provision of the *HBV Chronic Mouse* dataset, as well as Sébastien Baudart and Michaël Ska (GSK) for the data acquisition of the *ImmunoSenescence Human PBMC* dataset.

This preprint was created using the LaPreprint template (https://github.com/roaldarbol/lapreprint) by Mikkel Roald-Arbøl.

## Supplementary Materials

### Supplementary Tables

**Table S1.**
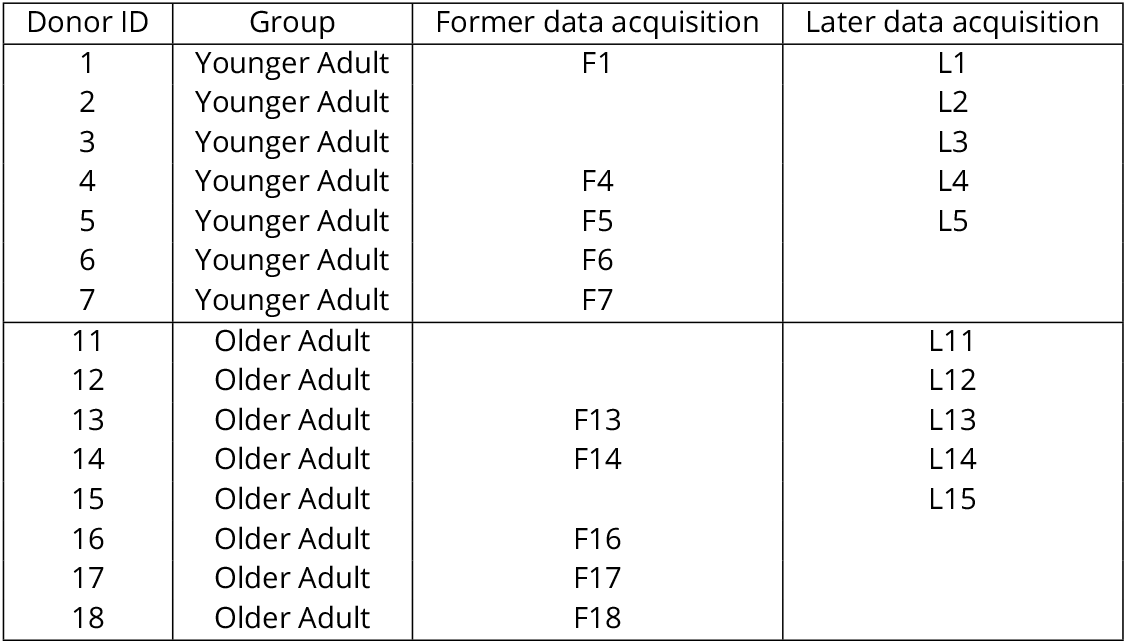
Sample groups and data acquisitions for the *ImmunoSenescence Human PBMC* dataset.

### Supplementary Figures

**Figure S1.**
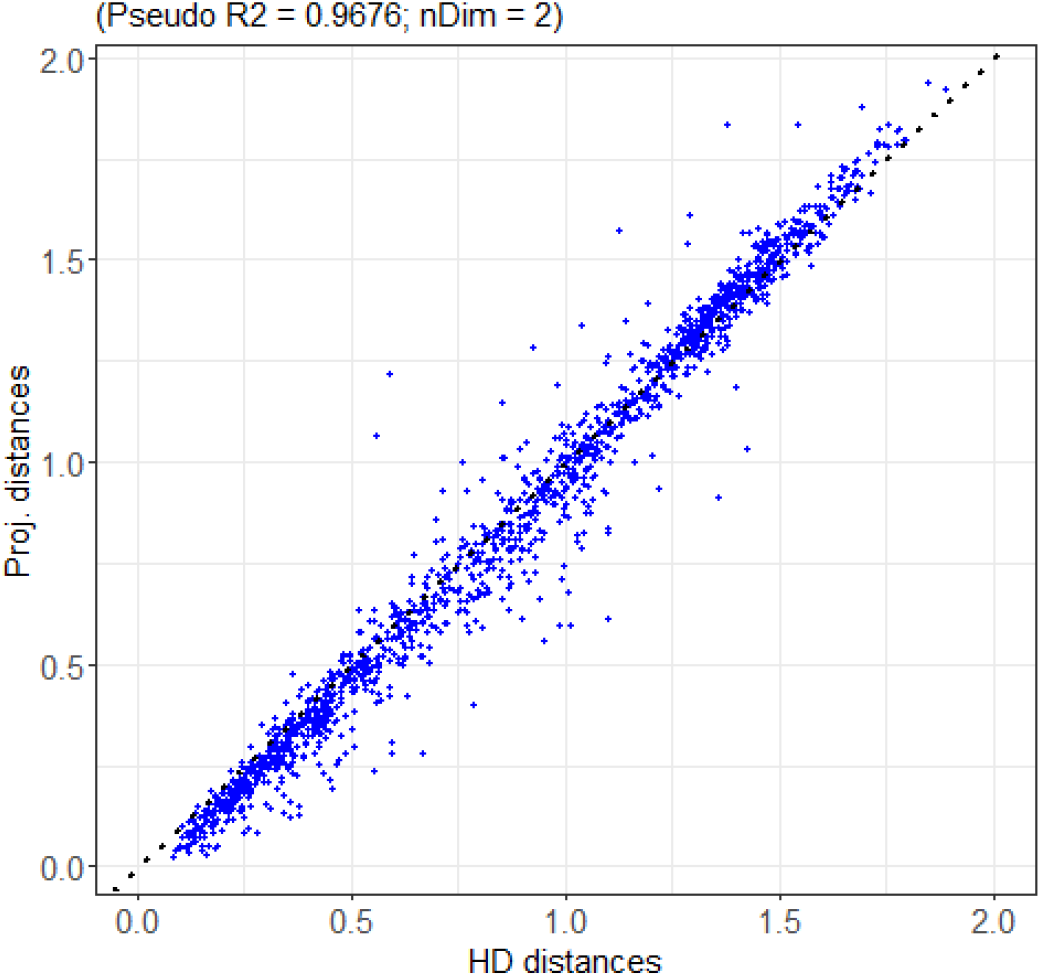
*Shepard’s diagram* for the projection of the *HBV Chronic Mouse* dataset. *CytoMDS* required 2 dimensions to reach the *pseudoR*^2^ quality threshold of 0.95. Each dot represents a sample pair, where the x value is the high dimensional *Earth Mover’s Distance* between the two samples and the y value is the obtained euclidian distance on the low dimensional projection. A projection of high quality results in most points being close to the ideal 45 degree identity line.

**Figure S2.**
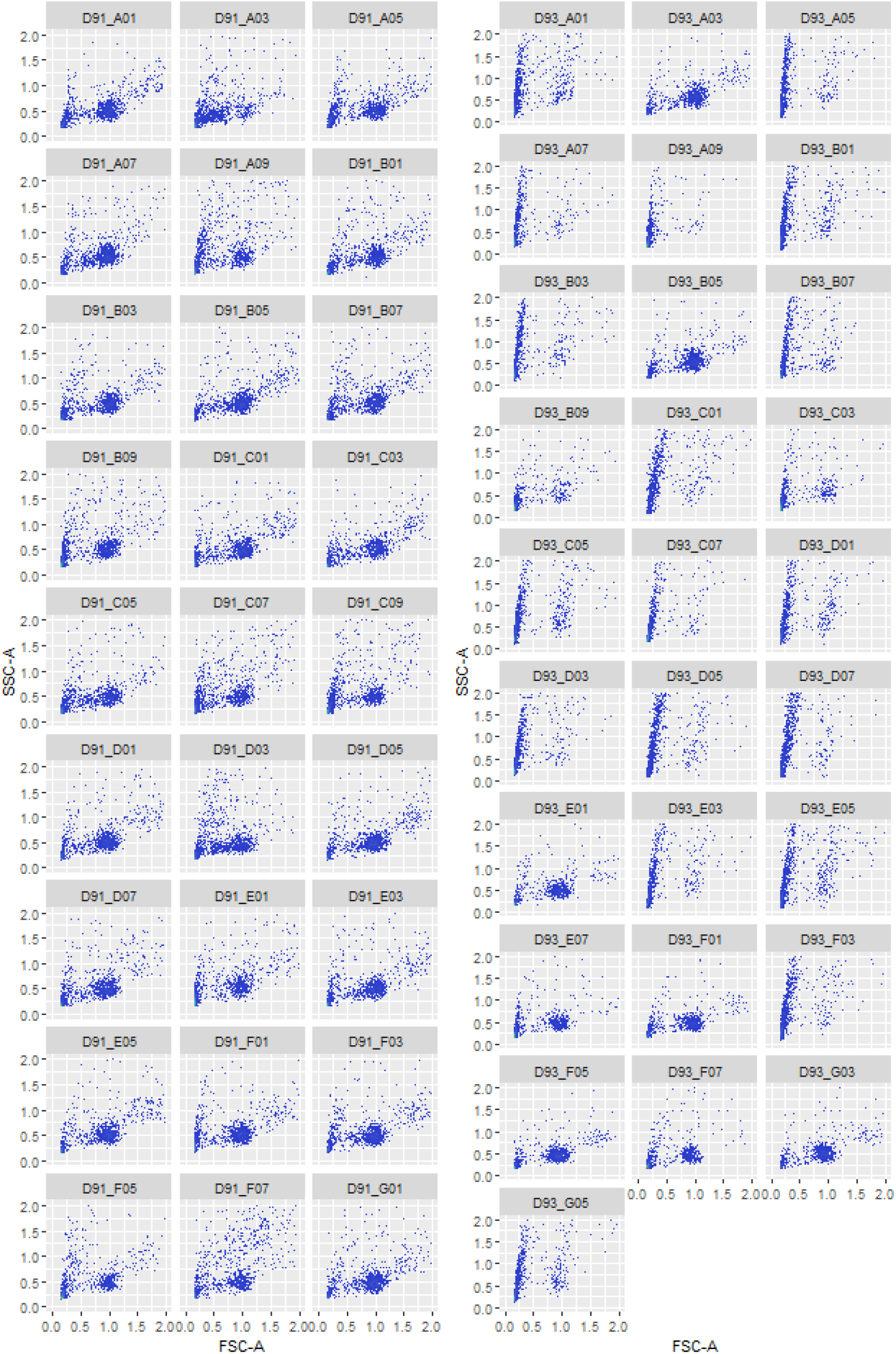
*FSC-A* vs. *SSC-A* 2D visualization of the 55 samples of the *HBV Chronic Mouse* dataset, revealing a clear difference of pattern between low quality samples and good quality samples, corresponding to the two sample groups in Figure 2. Compared to Figure 2, here the axes have undergone a linear mapping to the [0.0, 2.0] range, in order to keep the axis legends readable.

**Figure S3.**
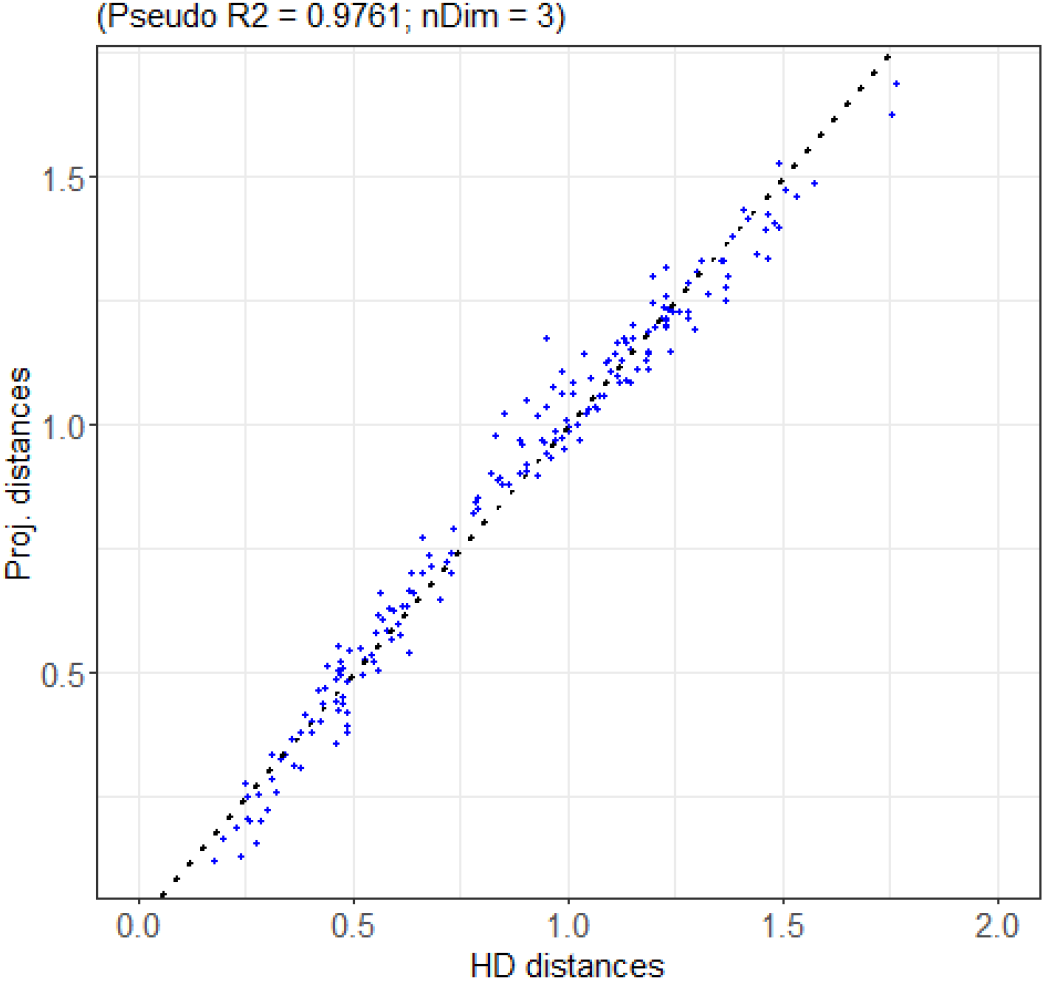
*Shepard’s diagram* for the projection of the *ImmunoSenescence Human PBMC* dataset. *CytoMDS* required 3 dimensions to reach the *pseudoR*^2^ quality threshold of 0.95. Each dot represents a sample pair, where the x value is the high dimensional *Earth Mover’s Distance* between the two samples and the y value is the obtained euclidian distance on the low dimensional projection. A projection of high quality results in most points being close to the ideal 45 degree identity line.

**Figure S4.**
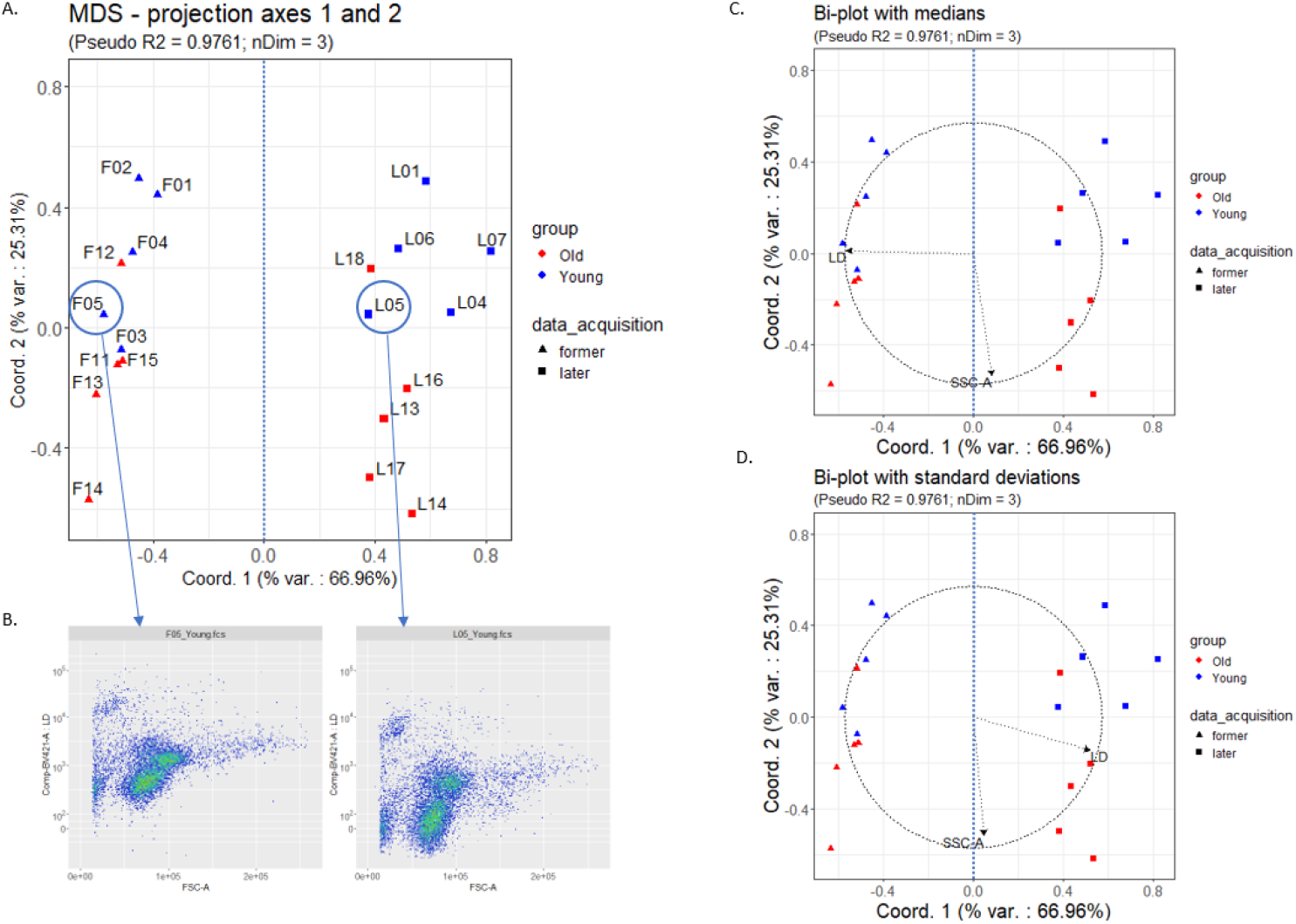
Low dimensional projection of the *ImmunoSenescence Human PBMC* dataset, 2D projection on axes 1 and 2. The projection plot (A) reveals a clear separation, along the first axis, between the samples of the former data acquisition (triangles), and the samples of the later data acquisition (squares). This is an obvious batch effect. According to the bi-plots using channel medians (C) and channel standard deviations (D), we can attribute this batch effect mainly to location and scale of *Live/Dead* channel. This is confirmed when plotting 2D representations of selected samples having opposite first coordinates (B).

**Figure S5.**
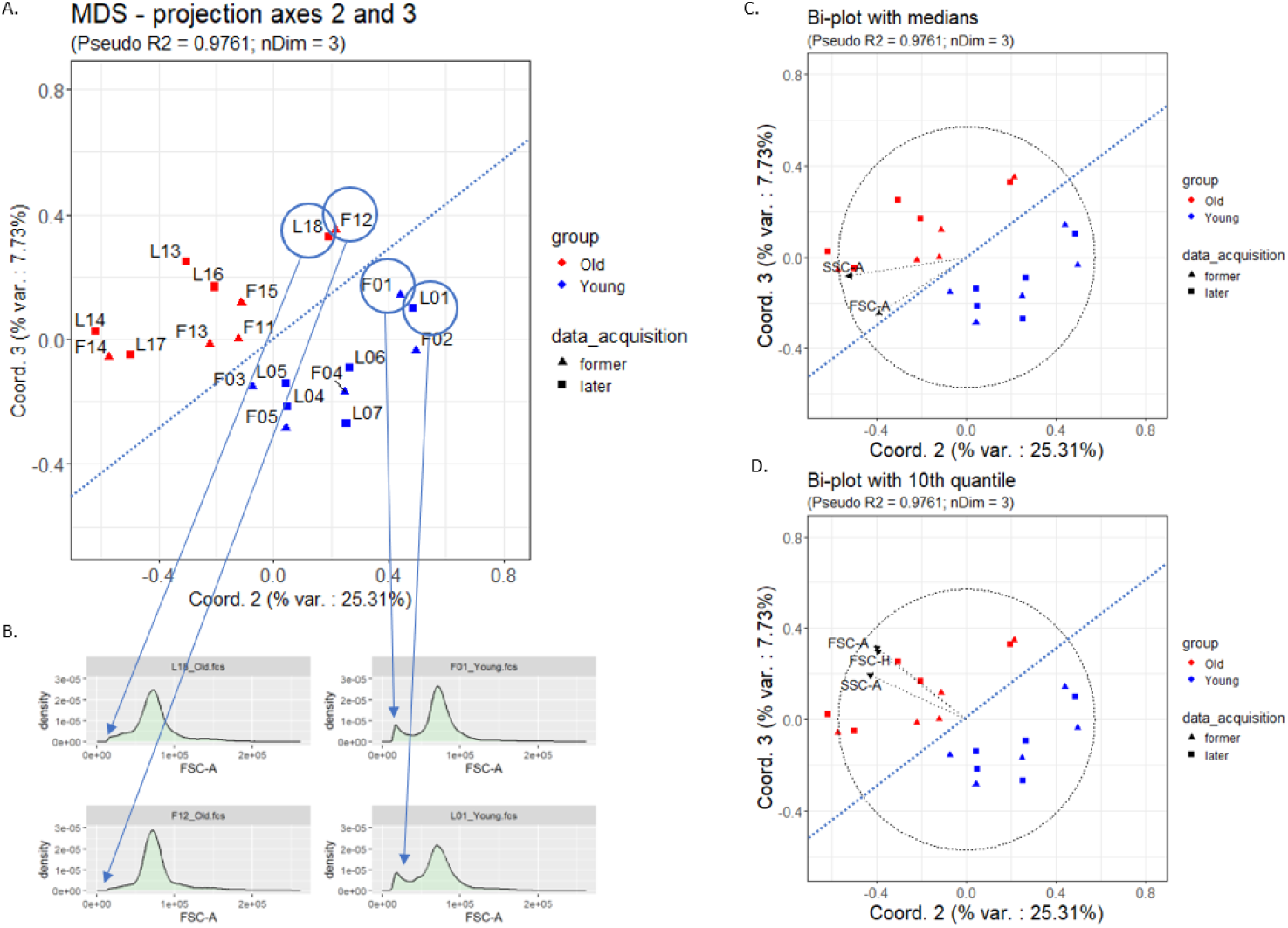
Low dimensional projection of the *ImmunoSenescence Human PBMC* dataset, 2D projection on axes 2 and 3. On the projection plot (A), the two groups (‘old’ in red vs. ‘young’ in blue), can be separated along a direction that is a linear combination of axes 2 and 3. Bi-plots using channel medians (C) does not highlight any channel median strongly correlated with the direction of separation. However, a bi-plot with a well-chosen statistics, i.e. the 10th percentile, enables the identification of a strong correlation with this quantile for the *FSC* and *SSC* channels. This can be confirmed when comparing the marginal distributions of the *FSC-A* channel for selected samples from the two sides of the border (B).

**Figure S6.**
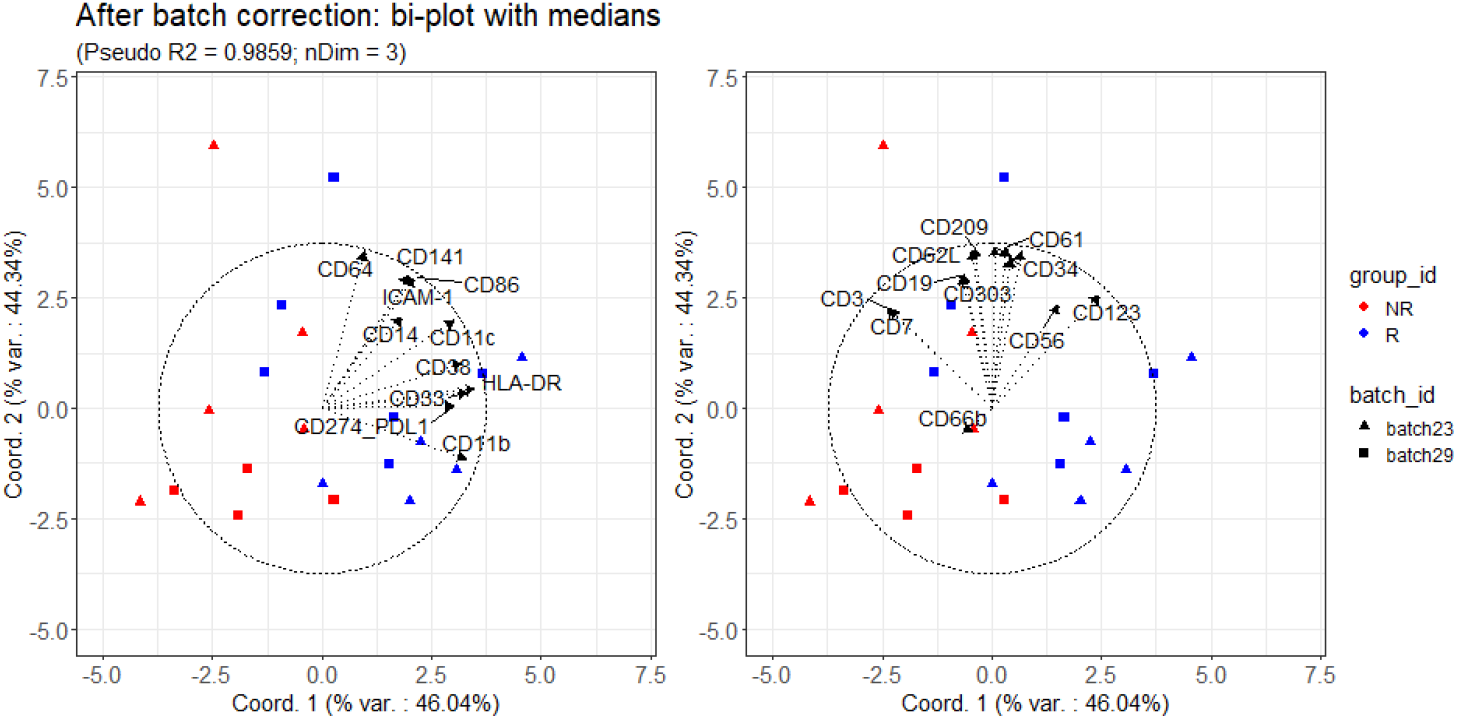
Bi-plots of the projection of the *Krieg_Anti_PD1* dataset on the first two axes, after batch correction, and representing all channel medians, split into two groups. In the original study (Krieg et al., 2018), it was highlighted that the frequency of a small subpopulation of CD14^+^ CD33^+^ HLA-DR^hi^ ICAM-1^+^ CD64^+^ CD141^+^ CD86^+^ CD11c^+^ CD38^+^ PD-L1^+^ CD11b^+^ monocytes in baseline samples was positively associated with responder status following immunotherapy treatment. On the left plot, we selected these 11 markers and on the right plot, the other 14 markers are represented for comparison. The intensity of the markers of the first group tends to be more correlated with the direction separating responders (blue) and non responders (red).

**Figure S7.**
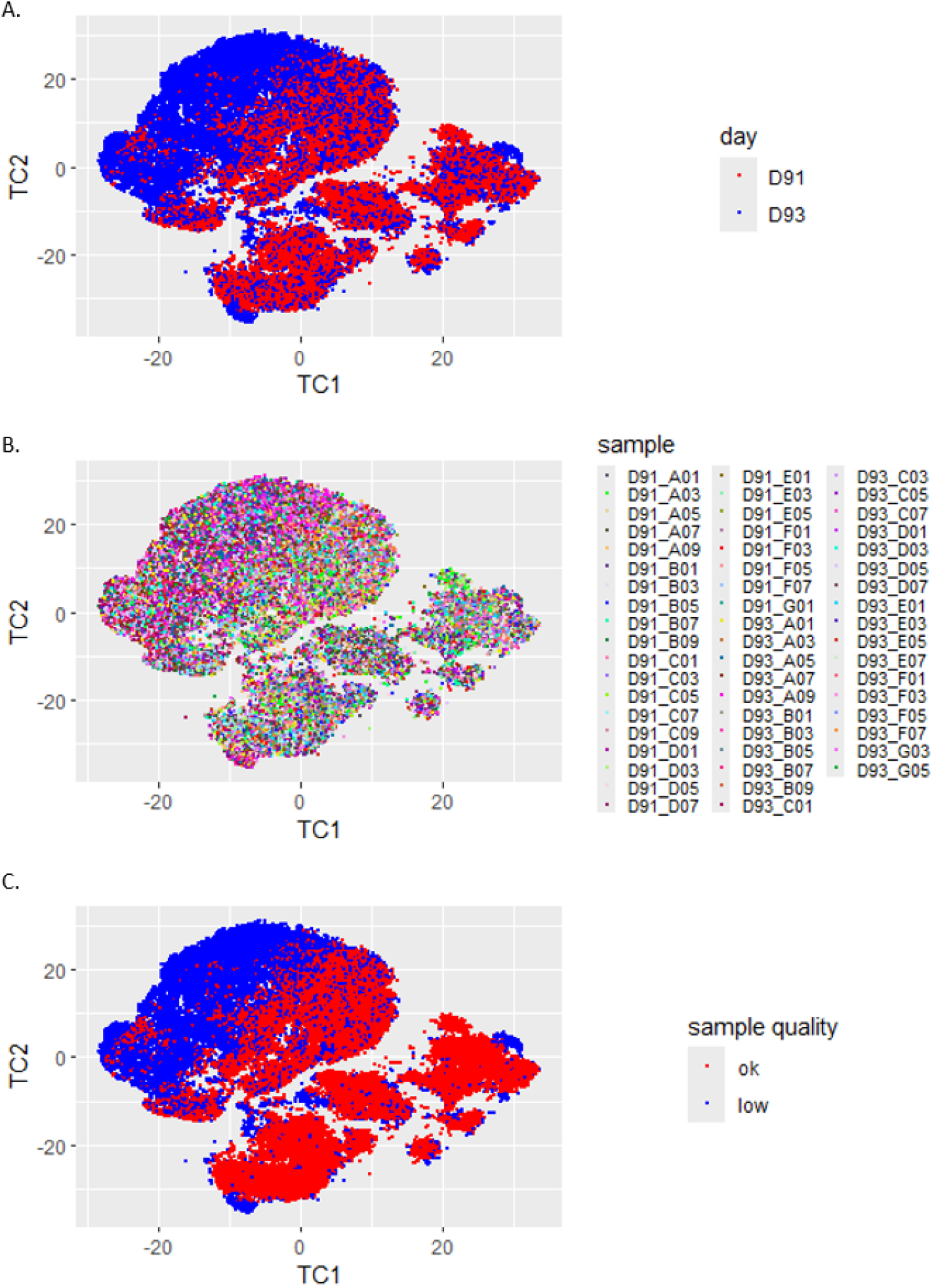
tSNE of the *HBV Chronic Mouse* dataset. Input data consist of a sub-sampling of 50,000 random events from the 55 samples, with all 12 markers included in the calculation. Prior to sub-sampling, channel intensities were transformed using respectively bi-exponential transformations for fluorescent channels, and linear transformations for *FSC/SSC*, as described in the Results section. Highlighting the displayed events by acquisition day (A) suggests the existence of a corresponding batch effect. However a substantial amount of the blue dots are mixed with the red ones, because the effect is linked to a quality issue affecting only a subset of D93 samples. Highlighting the displayed events by sample (B) is uninformative, due to the high number of samples, and high number of events per sample. (C) Colouring the events by quality cluster - which was identified thanks to *CytoMDS* analysis - seems to show less residual dots mixing than in (A).

**Figure S8.**
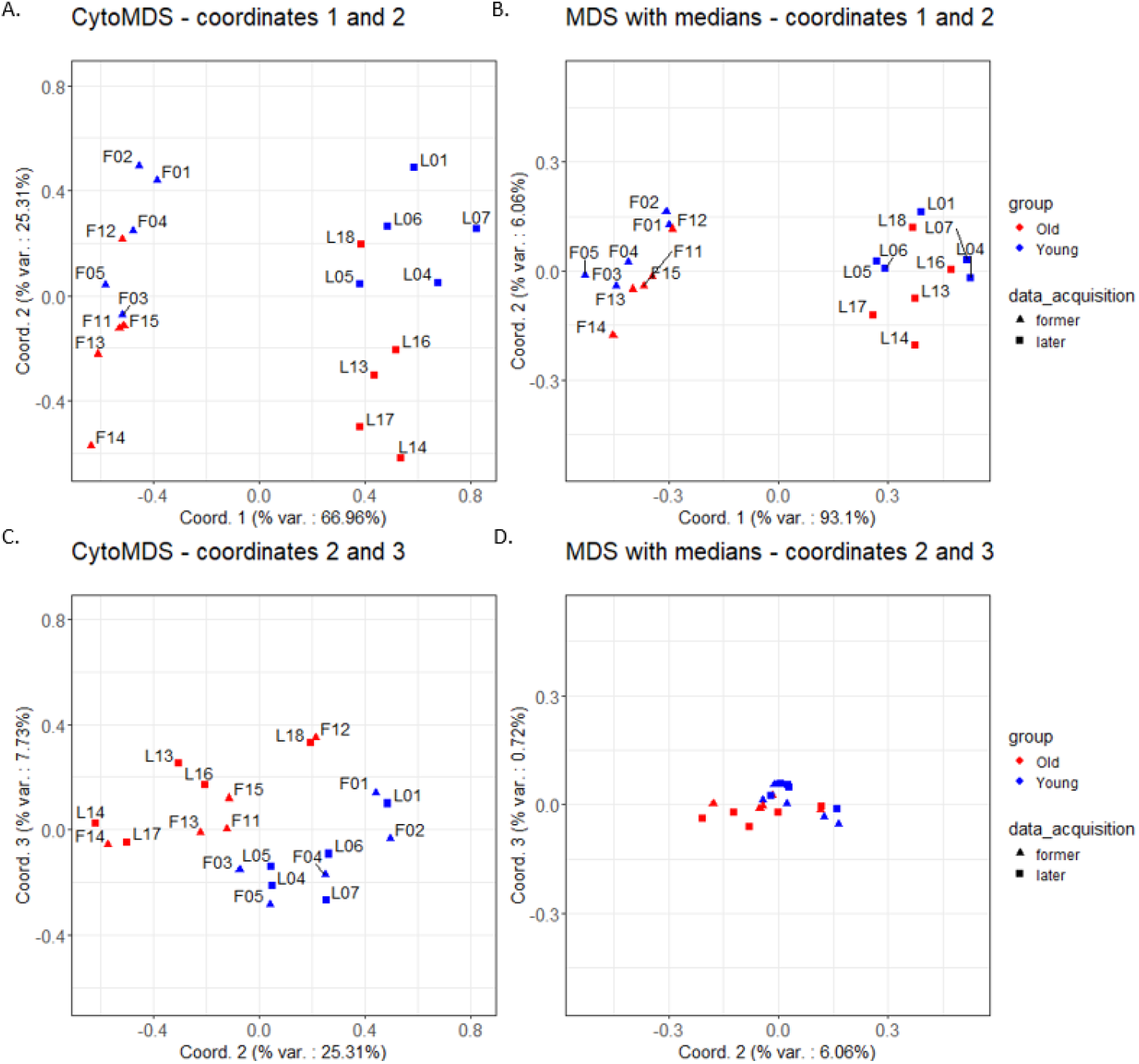
Low dimensional projections of the *ImmunoSenescence Human PBMC* dataset. The above plots show a comparison of the projection on axes 1 and 2 of respectively the *CytoMDS* method (A) - which uses *Earth Mover’s Distances* to assess dissimilarity between samples - and *MDS* using channel medians as input (B). The two plots show similar patterns. Similarly, plots (C) and (D) show the same comparison, now using axes 2 and 3. Here, *CytoMDS* projection (C) shows a visible separation between the two sample groups. However, on the right plot (D), there is little variability left between data points. This difference can be explained by the fact that the direction of separation between the two sample groups in the *CytoMDS* projection could be associated to the 10th quantile of the *FSC* and *SSC* channel distributions (Figure S5), but not to channel medians.

### Supplementary Results - benchmarking of CPU time for *EMD* computation

In this section, we present a benchmarking of the CPU time required to compute the *EMD* between two cytometry samples, as a function of number of markers and events. To perform this study, we randomly selected two cytometry samples from the *ImmunoSenescence Human PBMC* dataset, and we pre-processed these two samples by applying compensation and scale transformation, as described in the Results section. From these two pre-processed samples, we created two large simulated samples by sampling 10,000,000 events with replacement from the original sample event sets, while keeping the original 28 markers - 26 fluorescent markers and (FSC-A, SSC-A). Finally we added a gaussian random noise with a low standard deviation (0.05) to the scaled intensities.

With these two large simulated reference samples, we let *CytoMDS* calculate their *EMD*, which we considered their reference *EMD*. Then, we launched computational experiments by re-calculating the *EMD* for various combinations of randomly selected subsets of markers or events, and recorded the CPU time needed for each *EMD* run. All calculations were performed with a single core on a standard laptop computer with a 11th Gen Intel-Core i7 @ 2.80 Ghz CPU and 8GB of RAM. These computational experiments were repeated 20 times each. The results are summarized on Figures S9 and S10.

Figure S9 (A) shows that the calculation for the full reference samples with 10 million events - last box plot on the right - took slightly less than one minute to run. This reference time estimates an upper bound for a realistic *EMD* calculation time, since 10 million events and 28 markers is a reasonably large sample size in comparison with usual flow cytometry datasets. The plot also shows that the calculation time decreases steadily as a function of decreasing number of events, with a calculation time of less than one second for 100,000 events. For small samples with less than 1,000 events, a lower plateau is observed for the computation time. In Figure S9 (B), we investigate to what extend the *EMD* using a sub-sampling of a decreasing number of events, remains a good approximation of the true reference *EMD*. This plot shows that sub-sampling 100,000 events is more than enough to obtain an excellent approximation. This means that, even for large sample cases, when the user is not willing to wait up to one minute to obtain the *EMD*, they can still use sub-sampling to obtain satisfactory approximations with much shorter calculation times.

Finally, as is shown in Figure S10, the *EMD* calculation time evolves roughly as a linear function of number of markers. This is expected, since *CytoMDS* approximates the *EMD* by the sum of the per marker *EMD*s, each of them being calculated independently (see Methods section).

**Figure S9.**
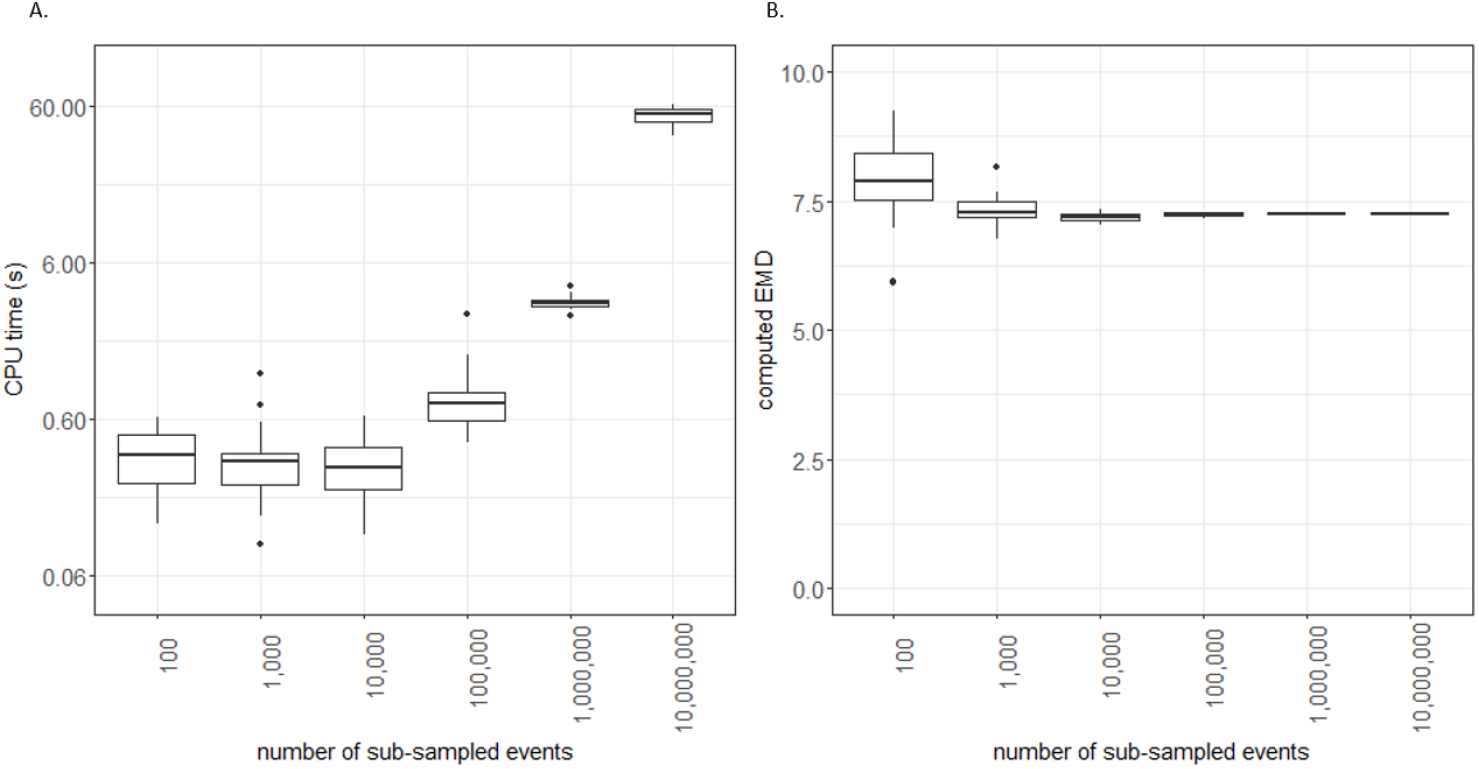
Distribution of calculation time (A) and obtained distance (B) when computing the *EMD* between the two reference samples, as a function of number of events. Each random sub-sampling of events and *EMD* calculation were run 20 times

**Figure S10.**
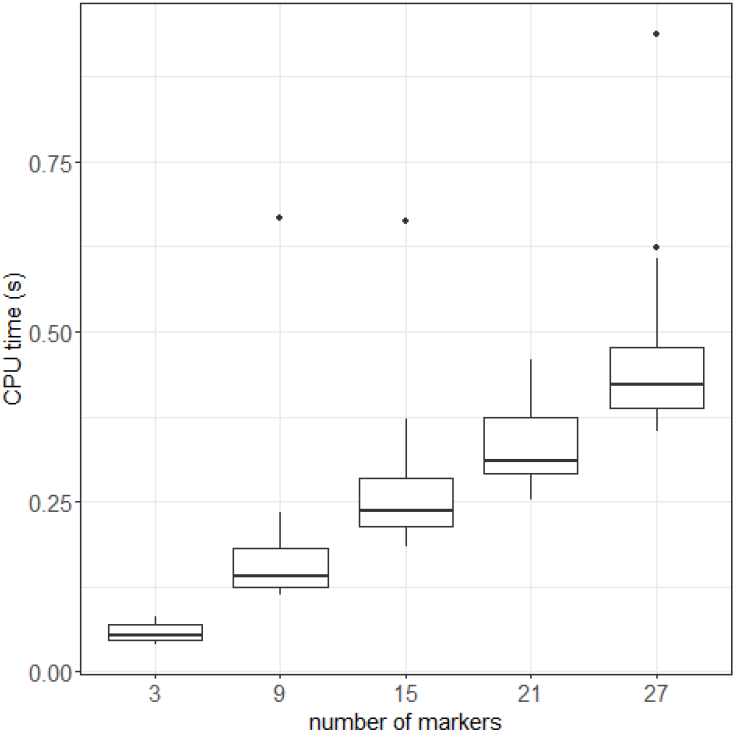
Distribution of calculation time when computing the *EMD* between the two reference samples, as a function of number of included marker, for a sub-sampling of 100,000 events. Each random marker selection and *EMD* calculation were run 20 times

